# A Machine Learning-based Framework to Identify Type 2 Diabetes through Electronic Health Records

**DOI:** 10.1101/078634

**Authors:** Tao Zheng, Wei Xie, Liling Xu, Xiaoying He, Ya Zhang, Mingrong You, Gong Yang, You Chen

## Abstract

**Objective:** To discover diverse genotype-phenotype associations affiliated with Type 2 Diabetes Mellitus (T2DM) via genome-wide association study (GWAS) and phenome-wide association study (PheWAS), more cases (T2DM subjects) and controls (subjects without T2DM) are required to be identified (e.g., via Electronic Health Records (EHR)). However, existing expert based identification algorithms often suffer in a low recall rate and could miss a large number of valuable samples under conservative filtering standards. The goal of this work is to develop a semi-automated framework based on machine learning as a pilot study to liberalize filtering criteria to improve recall rate with a keeping of low false positive rate.

**Materials and Methods:** We propose a data informed framework for identifying subjects with and without T2DM from EHR via feature engineering and machine learning. We evaluate and contrast the identification performance of widely-used machine learning models within our framework, including k-Nearest-Neighbors, Naïve Bayes, Decision Tree, Random Forest, Support Vector Machine and Logistic Regression. Our framework was conducted on 300 patient samples (161 cases, 60 controls and 79 unconfirmed subjects), randomly selected from 23,281 diabetes related cohort retrieved from a regional distributed EHR repository ranging from 2012 to 2014.

**Results:** We apply top-performing machine learning algorithms on the engineered features. We benchmark and contrast the accuracy, precision, AUC, sensitivity and specificity of classification models against the state-of-the-art expert algorithm for identification of T2DM subjects. Our results indicate that the framework achieved high identification performances (~0.98 in average AUC), which are much higher than the state-of-the-art algorithm (0.71 in AUC).

**Discussion:** Expert algorithm-based identification of T2DM subjects from EHR is often hampered by the high missing rates due to their conservative selection criteria. Our framework leverages machine learning and feature engineering to loosen such selection criteria to achieve a high identification rate of cases and controls.

**Conclusions:** Our proposed framework demonstrates a more accurate and efficient approach for identifying subjects with and without T2DM from EHR.

## Background and Significance

Type 2 diabetes mellitus (T2DM) is a major disease with high penetrance in humans around the globe, a trend that is still on the rise [1–2]. T2DM is a leading cause of morbidity and mortality and contributes to increased risks of heart disease by 2 to 4 times [1]. A significant number of research investigations have been devoted to it, notably by means of genome-wide association study (GWAS) and phenome-wide association study (PheWAS) in hope of detecting more associations between genotypes and phenotypes [3–10, 23–26, 36]. To discover diverse genotype-phenotype associations affiliated with T2DM via PheWAS and GWAS, more cases (subjects with T2DM) and controls (subjects without T2DM) are required to be identified from electronic health records (EHR) [11–12, 34–35].

A widely adopted approach for identifying subjects with and without T2DM is to have human experts (e.g., experienced physicians) manually design algorithms based on their experience and examination of EHR data [11, 13–15]. However, such strategies increasingly prove to be limited and not scalable [11, 13, 15] due to the laborious process of human intervention and rule abstraction capabilities of experts. Furthermore, expert algorithms are often designed with conservative identification strategy, thus may fail to identify complex (e.g., borderline) subjects and miss a significant number of potential T2DM cases and controls. In research settings such as GWAS and PheWAS, accumulating large sample sizes is often highly desirable and discarding valuable samples will influence the potentiality to discover diverse genotype-phenotype associations [26, 36]. A disease may be caused by the joint effects of multiple single nucleotide polymorphism (SNPs) (i.e. heterogeneity), while a SNP may lead to multiple diseases (i.e. pleiotropy) [32–34]. Involving more cases with diverse phenotypic characteristics such as comorbidities will enrich the association studies between phenotypes and genotypes. Given the limitations in high missing rate and laborious manual intervention, it is increasingly challenging for expert algorithms to scale to the ever-increasing volumes of diabetes related EHR data, secondary use and evolved GWAS and PheWAS studies [13, 15, 35].

Machine learning and data mining models are increasingly utilized in diabetes related research from EHR data (e.g., diabetes-related adverse drug effect, and association between periodontitis and T2DM) [27–29]. These studies have primarily focused on mining T2DM-related EHR data for clinical purposes, for instance, one such study aimed at forecasting clinical risk of diabetes from EHR [29]. The motivation and intended usage of the aforementioned work is different from ours, which aims to identify more cases and controls. Furthermore, the aforementioned study still has similar limitations in high missing rate [29]. To the best of our knowledge, very few studies have focused on reducing missing rate to identify more cases and controls for phenotyping purposes.

The goal of this work is to develop a semi-automated framework based on machine learning as a pilot study to identifying subjects with and without T2DM. Our method features two advancements: 1) low false positive rate; 2) high recall (i.e., detecting as many samples of interest as possible). To achieve these goals, we carefully approach feature engineering (i.e., construction of features for predictive modeling) by constructing representative features at three levels. We then train multiple popular machine learning models based on constructed features to identify cases and controls.

Our empirical evaluation is based on three years (ranging from 2012 to 2014) of EHR data from a large distributed EHR network consisting of multiple Chinese medical centers and hospitals in Shanghai, China. Our choice of this EHR repository is motivated by the fact that Chinese EHR data are often much worse than western EHR in terms of meaningful uses and data quality [18]. In addition, medical care in China often have non-standard unique procedures (such as wide adoption of traditional Chinese medicine) that are not represented in EHR and expert algorithms from elsewhere (such as from mainstream western counterparts), rendering standard or western expert algorithms less relevant. Given all such factors, the Chinese EHR repository provides an ideal test-bed for evaluating the accuracy and robustness of our proposed framework. In addition, the customization and empirical evaluation of a machine learning-based T2DM identification framework specifically for Chinese EHR is also of separate interest, which is under-explored despite constituting huge demand.

## Research Design and Methods

### Study Materials

Our investigations in this work focus on three years of EHR data (from Year 2012 to 2014). The data was stored in our centered repository, which has been managed by the District Bureau of Health in Changning, Shanghai since 2008. The EHR data generated from 10 local EHR systems are automatically deposited into the centralized repository hourly.

We have 123,241 patients in total within the investigated three years. We use a filtering strategy to pre-select patients as our candidate samples whose EHR data are related to diabetes. We pre-selected samples whose EHRs should satisfy at least one of the three criteria: i) diabetes related diagnosis, ii) diabetes related medication and iii) diabetic laboratory test. Through this process, we managed to obtain 23,281 patient samples with diabetes related information. Our data preparation workflow is summarized in Figure 1.

**Figure 1.**
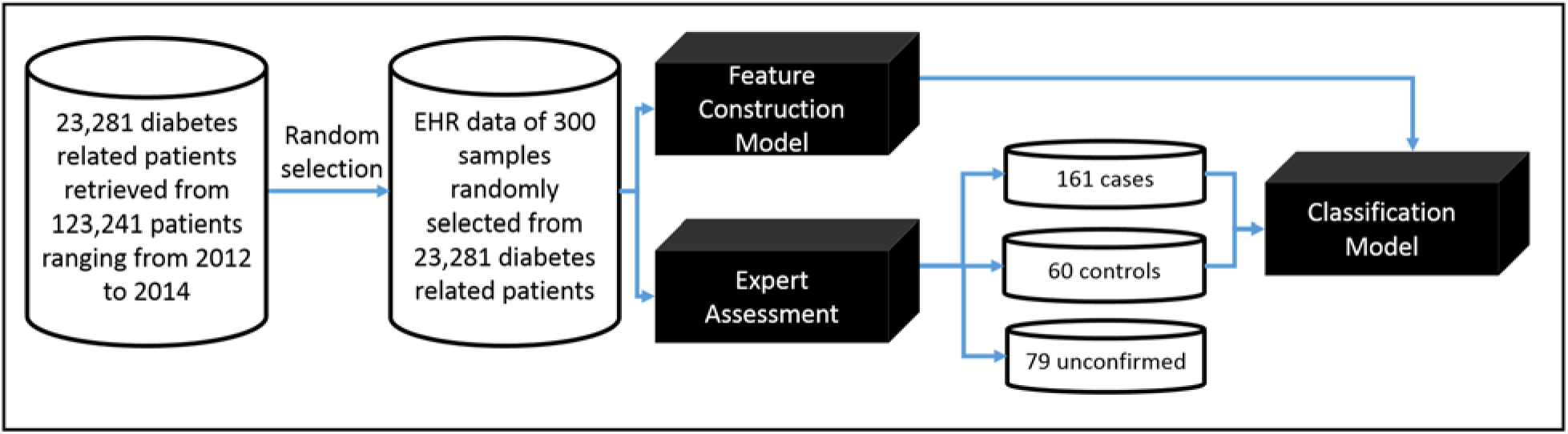
A machine learning-based framework to identify subjects with and without T2DM from EHR data. The *case* refers to subjects with T2DM, and *control* refers to non-T2DM subjects.

Our framework is based on supervised learning (e.g., classification, to be specific), which requires labeled training samples. Thus, we invited two clinical experts experienced in diabetes to assess EHRs of samples and label these samples into three categories: case, control and unconfirmed. We point out that, as is common for similar efforts, our expert review process is based on manually judging the whole record of each patient instead of only considering the selected few criteria in our data filtering or baseline expert algorithms (introduced later). Due to huge amount of manual effort in the expert reviewing process, as a pilot study, we randomly selected 300 samples out of the 23,281 pre-selected ones and concentrated our reviewing efforts on the smaller subset. For the investigated 300 selected samples, there are 20,384 records (e.g., diagnostic notes, communication notes and summary notes). Samples with two confirmed labels of T2DM from both clinicians will be considered as cases, samples with two confirmations of Non-T2DM considered as controls. The other samples with conflicting labels or two confirmations of un-determined from two clinicians will be denoted as unconfirmed ones. Through clinicians’ assessments, we obtained 161 cases, 60 controls and 79 unconfirmed samples. For double check, we noticed that of the unconfirmed 79 samples, most (78.3%) are severely incomplete in their EHR documentation, which are not be suitable for EHR-based phenotyping. In order to reduce negative influences of incomplete EHRs on performances of our classification models, we dropped 79 unconfirmed samples.

Through our assessing processes of cases and controls, the separation range between cases and controls in our study is narrower than that in traditional expert algorithms as shown in Figure 2. This is because, in our study, controls refer to samples satisfying at least one of the following three criteria: i) one time of abnormal lab tests (HbA1C ≥6.0% or fasting plasma glucose ≥126 mg/dl or 2-hours plasma glucose≥200 mg/dl or random plasma glucose ≥200 mg/dl), ii) one time of prescribed diabetic medicine, and iii) one time of diabetic diagnosis. However, these controls were excluded in expert algorithms [11, 13, 15, 30]. However, the widely used expert algorithms selected controls whose EHR data should not include any of the three above mentioned diabetic related information. The selection criteria of expert algorithms will miss many controls. For instance, we investigated a number of control samples, who had high values of HbA1C (≥6.0%) recorded, but their fasting and post-meal blood sugars were normal. Another example is we found several controls whose records contained prescriptions of diabetic medications, but no diabetic diagnoses and laboratory tests were found in their records. One of reasons is the medications they prescribed were not for themselves, but for their friends or someone else.

**Figure 2.**
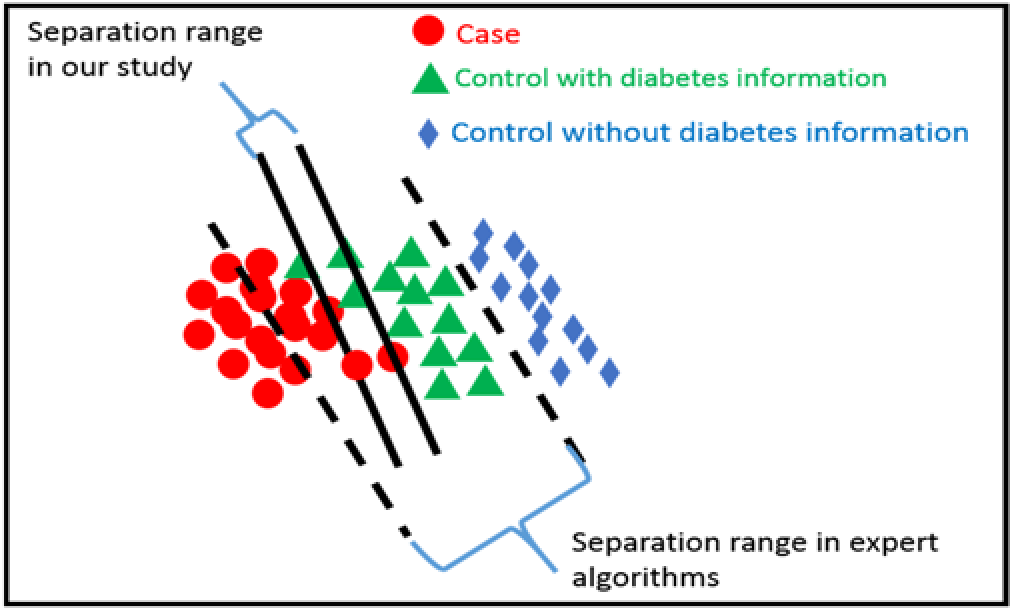
Separated boundary lines between cases and controls in our study and in traditional T2DM identification studies.

For the cases selection, expert algorithms selected samples whose EHR data should at least satisfy 2 of the following three requirements.

1. Abnormal laboratory tests (glucose ≥110 mg/dl or HbA1c ≥6.0%)
2. Diabetic medication
3. Diabetic diagnosis

Such selection process does not consider patients satisfying no more than 1 of the above three requirements but considering as T2DM patients through their related support information such as diabetic complications and self-reported body weight loss, persistent hunger, polyuria, and polydipsia. As a result, these cases were missed. According to our selection criteria, the range of separations between cases and controls are much smaller than expert algorithms as shown in Figure 2. We applied expert algorithms and our proposed framework to identify cases and controls in the same separation range (the range between two solid lines as shown in Figure 2). Both types of algorithms were studied on the same sources of 300 samples.

Our proposed framework includes feature construction, and classification models. Feature construction transforms raw EHR data into statistical features, which are further served as input entities to feed classification models (as shown in Figure 1). The expert algorithms extracted their three major features (abnormal laboratory tests, diabetic medication and diabetic diagnosis) in the same raw EHR data and then use their standards (workflows as depicted in the Figure A1 of Appendix F) to identify cases and controls.

### Feature construction

Constructing good features from EHR is often a must to warranty good prediction performance either for expert algorithms or machine learning-based models. This is because raw EHR data are often noisy, sparse, and contain unstructured information (e.g., text) that are not directly “computable”. Traditional researches on identifying subjects with and without T2DM were using selection strategies built on three features: diabetic diagnosis, diabetic laboratory tests and diabetic medications extracted from EHRs of investigated samples [11, 16]. Such researches are limited due to their high missing rates on identification of cases and controls. This is because such strategies applied a conservative selection criteria on cases and controls (e.g., satisfying two of the aforementioned three features) and were tested in a broader separation range between cases and controls (the range between two dashed lines as shown in Figure 2).

In our work, we include borderline (samples between two dashed lines of Figure 2), which can help to identify more cases and controls than traditional studies. To make the case/control identification more accurate, we need to incorporate more features than traditionally used. For instance, we constructed additional T2DM features such as self-reported diabetes related symptoms, and diabetic complications, and so on, in hope of better identifying borderline or more ambiguous samples. In total, we derived 110 features from seven sources (we denote this as First-Level features): “*demographic information*”, “*communication report*”, “*outpatient diagnosis report*”, “*inpatient diagnosis report*”, “*inpatient discharge summary*”, “*prescription report*” and “*laboratory test report*”, as summarized in Table 1 and with in-depth explanation off each feature in Table A1 of Appendix A.

**Table 1.** First-level Features constructed from source “*demographic information*”, “*communication reports*”, “*outpatients diagnosis reports*”, “*inpatients diagnosis reports*”, “*inpatients discharge summaries*”, “*prescription reports*” and “*laboratory test reports*”.

| Source category |  | Feature |
| --- | --- | --- |
| Demographic information |  | De-identification ID, age and gender of a subject. (feature ranging from f1 to f3 as shown in Table A1 of Appendix A) |
| Communication report | Self-reporting note | Number of times a subject reporting body weight loss, persistent hunger, polyuria, polydipsia, prescribed diabetes medicine or returning visits for diabetes in communication report. (feature ranging from f4 to f9 as shown in Table A1 of Appendix A) |
|  | Diagnosis code | Number of times codes of type 2 diabetes, diabetic retinopathy, diabetic neuropathy, diabetic eye disease, diabetic kidney disease, diabetic cerebral vascular disease or diabetic peripheral circulation appeared in communication report. (feature ranging from f10 to f17 as shown in Table A1 of Appendix A) |
|  | Diagnosis note | Number of times communication report containing notes of type 2 diabetes, diabetic retinopathy, diabetic neuropathy, diabetic eye disease, diabetic kidney disease, diabetic cerebral vascular disease or diabetic peripheral circulation disease. (feature ranging from f18 to f25 as shown in Table A1 of Appendix A) |
| Outpatient diagnosis record | Diagnosis code | Number of times codes of type 2 diabetes, diabetic retinopathy, diabetic neuropathy, diabetic eye disease, diabetic kidney disease, diabetic cerebral vascular disease or diabetic peripheral circulation were appeared in outpatient diagnosis record. (feature ranging from f26 to f33 as shown in Table A1 of Appendix A) |
|  | Diagnosis note | Number of times outpatient diagnosis record containing notes of type 2 diabetes, diabetic retinopathy, diabetic neuropathy, diabetic eye disease, diabetic kidney disease, diabetic cerebral vascular disease or diabetic peripheral circulation disease. (feature ranging from f34 to f41 as shown in Table A1 of Appendix A) |
| Inpatient discharge summary | Diagnosis notes | Number of times inpatient discharge summary containing notes of type 2 diabetes, diabetic retinopathy, diabetic neuropathy, diabetic eye disease, diabetic kidney disease, diabetic cerebral vascular disease or diabetic peripheral circulation disease. (feature ranging from f42 to f49 as shown in Table A1 of Appendix A) |
| Inpatient diagnosis record | Diagnosis codes | Number of times codes of type 2 diabetes, diabetic retinopathy, diabetic neuropathy, diabetic eye disease, diabetic kidney disease, diabetic cerebral vascular disease or diabetic peripheral circulation were appeared in inpatient diagnosis record. (feature ranging from f50 to f57 as shown in Table A1 of Appendix A) |
|  | Diagnosis notes | Number of times inpatient diagnosis record containing notes of type 2 diabetes, diabetic retinopathy, diabetic neuropathy, diabetic eye disease, diabetic kidney disease, diabetic cerebral vascular disease or diabetic peripheral circulation disease. (feature ranging from f58 to f65 as shown in Table A1 of Appendix A) |
| Prescription record | Medication | Number of prescriptions appearing in prescription report for oral hypoglycemic, insulin, Chinese traditional hypoglycemic, a mixture of western and Chinese traditional oral hypoglycemic, Epalrestat, Alpha-glucosidase inhibitor, Dipeptidyl peptidase IV(DPP-IV) inhibitors, Meglitinides, Sulfonylureas, Thiazolidinedione, Biguanides, Incretin Mimetics, GLP-1 (glucagon-like peptide 1) mimetics, compounds of sulfonylurea and thiazolidinedione, compounds of Biguanides and Dipeptidyl peptidase IV(DPP-IV) inhibitors, compounds of Biguanides and Sulfonylureas, or compounds of Biguanides and Thiazolidinedione. (feature ranging from f66 to f82 as shown in Table A1 of Appendix A) |
| Laboratory test report | Venous plasma glucose test | Number of times for 2-hours venous plasma glucose test, 2-hours venous plasma glucose test≥11.1mmol/l (200mg/dl), fasting venous plasma glucose test, fasting venous plasma glucose test ranging from 6.1 to 7.0 mmol/l (110 and 126 mg/dl), random venous plasma glucose test, or random venous plasma glucose test≥11.1mmol/l (200mg/dl). (feature f83, f84, f87, f88, f91, and f92 as shown in Table A1 of Appendix A) |
|  |  | The maximum value of 2-hours venous plasma glucose test, fasting venous plasma glucose test, or random venous plasma glucose test. (feature f85, f89, and f93 as shown in Table A1 of Appendix A) |
|  |  | The minimum value of 2-hours venous plasma glucose test, fasting venous plasma glucose test, or random venous plasma glucose test. (feature f86, f90, and f94 as shown in Table A1 of Appendix A) |
| | Peripheral plasma glucose test | Number of times for 2-hours peripheral plasma glucose test, 2-hours peripheral plasma glucose test≥11.1mmol/l (200mg/dl), fasting peripheral plasma glucose test, fasting peripheral plasma glucose test ranging from 6.1 to 7.0 mmol/l (110 and 126 mg/dl), random peripheral plasma glucose test, or random peripheral plasma glucose test≥11.1mmol/l (200mg/dl). (feature f95, f96, f99, f100, f103, and f104 as shown in Table A1 of Appendix A) |
|  |  | The maximum value of 2-hours peripheral plasma glucose test, fasting peripheral plasma glucose test, or random peripheral plasma glucose test. (feature f97, f101, and f105 as shown in Table A1 of Appendix A) |
|  |  | The minimum value of 2-hours peripheral plasma glucose test, fasting peripheral plasma glucose test, or random peripheral plasma glucose test. (feature f98, f102, and f106 as shown in Table A1 of Appendix A) |
| | HbA1C test | Number of times for HbA1c test, HbA1C test $\geq 6.5\%$. (feature f107, and f108 as shown in Table A1 of Appendix A) |
|  |  | The maximum value of HbA1C test (feature f109 as shown in Table A1 of Appendix A) |
|  |  | The minimum value of HbA1C test (feature f110 as shown in Table A1 of Appendix A) |

Notably, the features includes supporting materials for T2DM such as diabetic complications (e.g., diabetic retinopathy, diabetic neuropathy, diabetic cerebral vascular and diabetic peripheral circulation diseases), self-reported symptoms (e.g., self-report of body weight loss, persistent hunger, polyuria, and polydipsia), additional Chinese traditional medications and more laboratory test items (e.g., two-hours, fast and random glucose tests).

For the features in the medication category, we list investigated medicines related with T2DM treatments as in Table A2 of Appendix B. Notably, to tailor for Chinese EHR, we added additional Chinese traditional medicine, and mixtures of Chinese traditional and western medicine into the medication list. This is due to observation that T2DM patients are usually treated with a combination of Chinese traditional and western medicine in China, which is different from the common practice (i.e., western medicine only) of the western world and was thus neglected by western EHR-oriented studies.

For the diagnosis notes related features, we use regular expressions combining positive notes and negative notes as depicted in Table A3 of Appendix C to build each of them.

### Feature summarization

Features (as shown in Table A1 of Appendix A) cover seven EHR sources, however, some sources have the same type of features. For instance, f_10_ in the source of “*communication report*”, f_26_ in “*outpatient diagnosis record*”, and f_50_ in “*inpatient diagnosis record*” have the same definition on the counting of diagnosis codes. These features are highly correlated with each other, which will influence performances of computational models to do classification [17, 19, 22]. And thus we merge correlated features into one feature by summarizing them. For instance, f_10_, f_26_ and f_50_ are summarized as a new feature f’_10_= f_10_+f_26_+f_50_, which represents the total number of times T2DM diagnosis codes appearing in “*communication reports*” (f_10_), “*outpatient diagnose records*” (f_26_) and “*inpatient diagnose records*” (f_50_) respectively. By using the same way, we summarize all similar features across the seven sources into 36 features as shown in Table A4 of Appendix D. At the same time, the features within a source are also correlated, so we transform 36 features into final 8 features through summarizing correlated features within a source. The final 8 features are listed in Table A5 of Appendix E.

### Classification

We use several widely-used classification model such as k-Nearest-Neighbors (kNN), Naïve Bayes (NB), Decision Tree (J48), Random Forest (RF), Support Vector Machine (SVM) and Logistic Regression (LR) to model patterns of cases and controls based on our extracted features and then use the models to test the ability of our extracted features on identifications of T2DM subjects. These classification models are frequently utilized in a wide range of fields, and are recognized as popular choices for classification tasks [20–21, 37].

## Results

### Experimental set-up

Our framework adopts feature engineering by abstracting the EHR data at three different levels. This ensures to leverage more available data while maximizing predictive power. For the 221 T2DM samples (160 cases and 61 controls) collected with expert labels, we first construct 110 features (as mentioned before; see also Table A1 in Appendix A) to represent their EHR data. This is roughly a summarized and structured version of the raw EHR data. To prevent data sparsity and noise, we then derive higher-level features by condensing the data into 36 features (see Table A4 in Appendix D) and 8 features (see Table A5 in Appendix E), respectively. Such abstraction is mainly based on common knowledge of EHR data hierarchies.

In our framework, we apply several widely-used machine learning models, including kNN, NB, J48, RF, SVM and LR. The goal is to find out the comparative performance of machine learning models against expert algorithms. We used Weka package to apply these models on our engineered features [31]. We perform training and evaluation on different abstraction levels of feature sets, e.g., on the 107 aforementioned features^1^ (the first level of features; as shown in Table A1 of Appendix A), 33 features (the second level of features; as shown in Table A4 of Appendix D), and 5 features (the third level of features; as shown in Table A5 of Appendix E), respectively. We conduct extensive comparison of different classifiers on the same level of features, as well as performance across the three different levels of feature sets described before. Furthermore, we use the state-of-the-art expert algorithm [11] as a benchmark baseline, which is widely adopted by several large EHR and genetics consortia studies. We emphasize that the expert algorithm [11] is evaluated on the same raw EHR data as mentioned before.

We also point out that our primary focus of this work is to demonstrate feasibility/suitability of machine learning-based framework for the given task, and to provide general model recommendations and suggestions. Comprehensive and systematic benchmark of different machine learning models is not the main focus and is a separate topic with extensive literature. To keep our work focused and data-efficient, we adopt default recommended model parameters instead of performing hyperparameter tuning, since the latter often requires setting aside independent validation datasets, which may not be a wise option given our relatively small (and valuable) expert-labeled dataset. Our decision thresholds in certain models are also based on default configurations in Weka software [31]. For instance, in logistic regression, we use p=0.50 as the classification cut-off.

### Performance of classification models

For each classifier and each level of feature set, we conduct 4-fold cross-validation and report on the average performance and standard deviation. We demonstrate the prediction accuracy results in Figure 3, which measures the ratio of correctly predicted samples. In Figure 4, the prediction sensitivity (also called recall) results are reported, which measures the ratio of true positives against all positives. Lastly, in Figure 5, we plot the specificity, which denotes the proportion of true negatives of all negatives. The precision (or positive predictive value) results are illustrated in Figure 6. For more comprehensive comparison, we also present the area under the receiver operating characteristic (ROC) curve (AUC) in Figure 7, which demonstrates the trade-off between false positive and true positive rates (larger AUC generally implies better performance). All detailed numerical metrics are also summarized in Table 2.

**Figure 3.**
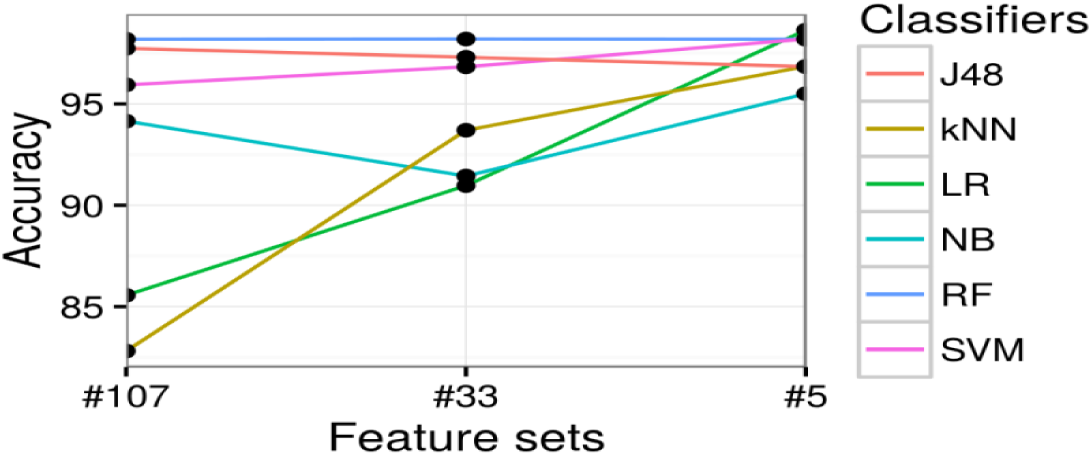
Prediction accuracy (y-axis) with different feature sets (x-axis), categorized by different classifiers (different lines plotted).

**Figure 4.**
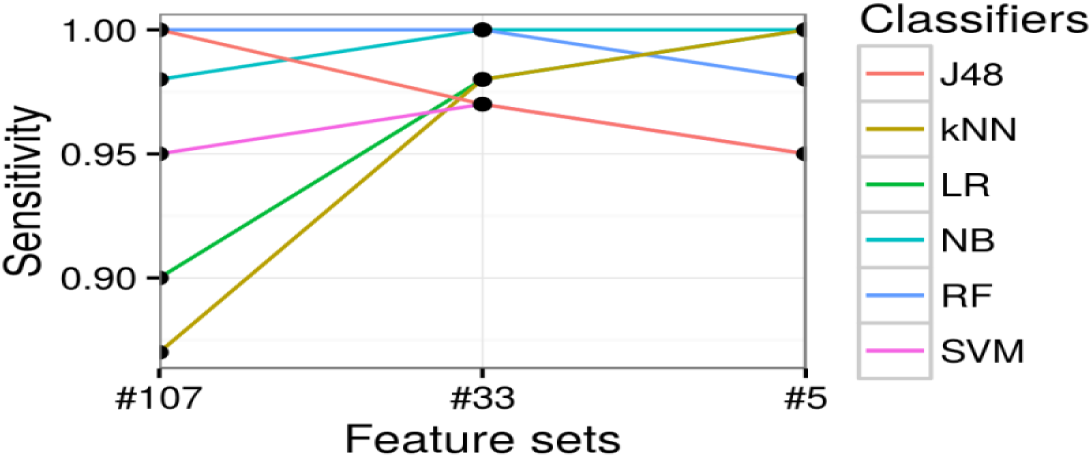
Prediction sensitivity [True positive rate] (y-axis) with different feature sets (x-axis), categorized by different classifiers (different lines plotted).

**Figure 5.**
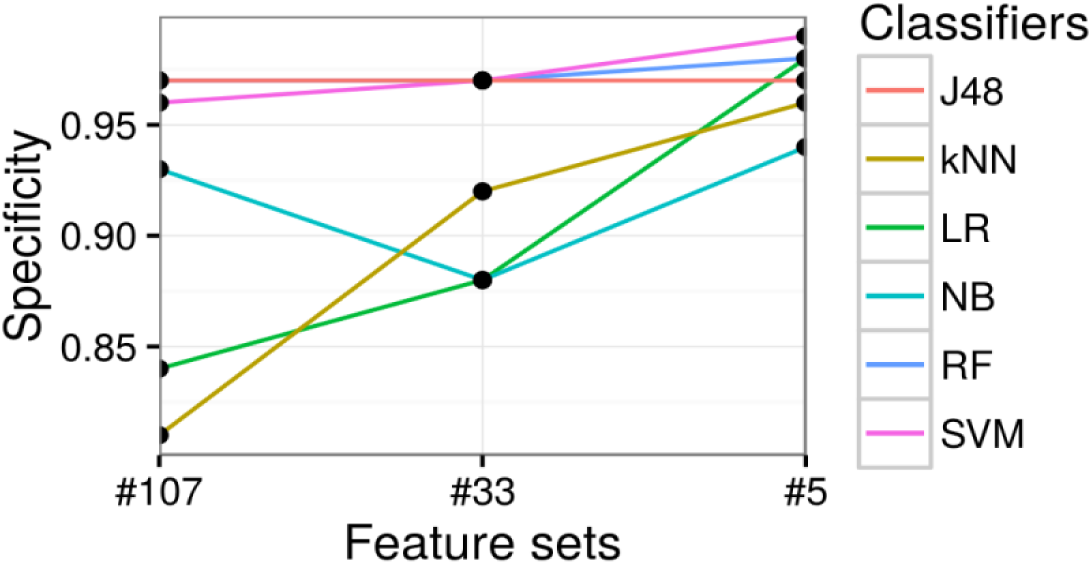
Prediction specificity [True negative rate] (y-axis) with different feature sets (x-axis), categorized by different classifiers (different lines plotted).

**Figure 6.**
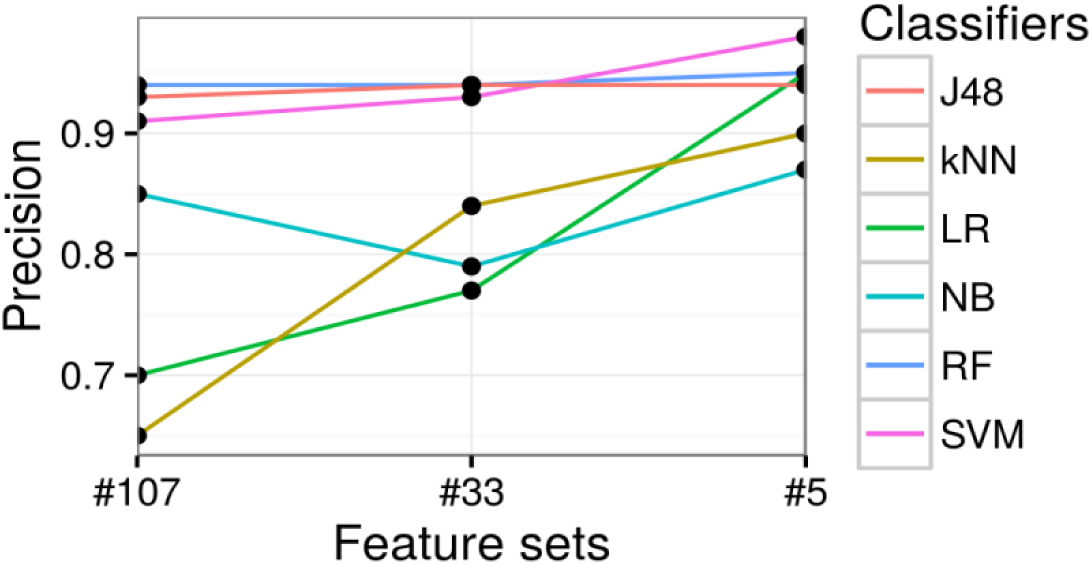
Prediction precision [Positive predictive value] (y-axis) with different feature sets (x-axis), categorized by different classifiers (different lines plotted).

**Figure 7.**
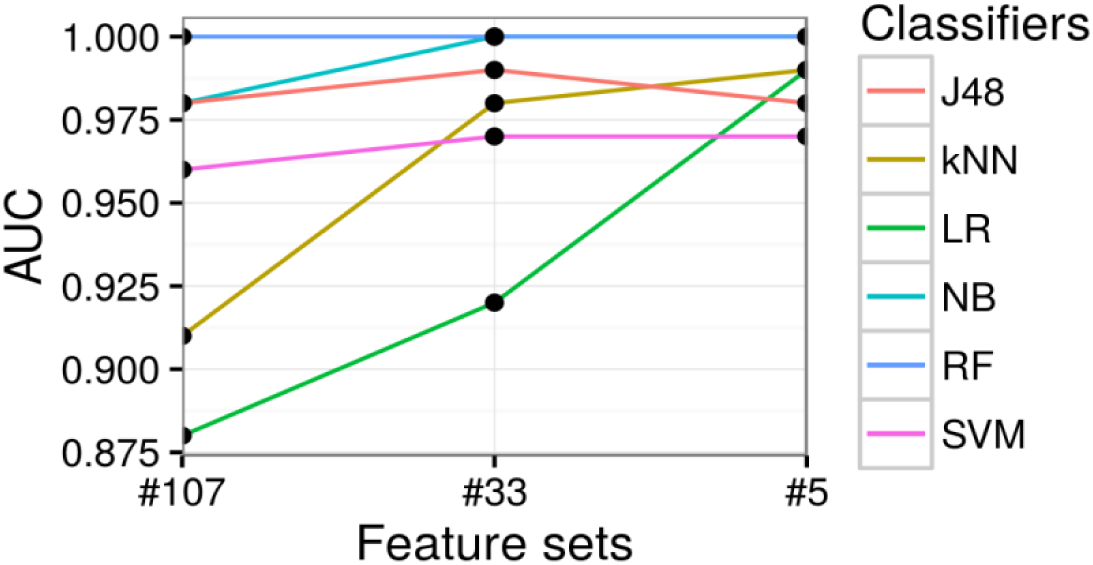
Prediction AUC (y-axis) with different feature sets (x-axis), categorized by different classifiers (different lines plotted).

**Table 2.** Comparison of different classifiers and the expert algorithm (baseline), measured by their average performance (and standard deviation) in cross-validation.

| Classifiers | Feature Sets | Accuracy | Sensitivity | Specificity | Precision | AUC |
| --- | --- | --- | --- | --- | --- | --- |
| Expert Algorithm | --- | 0.84 | 0.78 | 1.00 | 1.00 | 0.71 |
| LR | #107 | 0.86 (0.06) | 0.90 (0.09) | 0.84 (0.10) | 0.70 (0.11) | 0.88 (0.07) |
|  | #33 | 0.91 (0.04) | 0.98 (0.03) | 0.88 (0.06) | 0.77 (0.07) | 0.92 (0.03) |
|  | **#5** | **0.99 (0.01)** | **1.00 (0)** | 0.98 (0.01) | 0.95 (0.03) | 0.99 (0.01) |
| **NB** | **#107** | 0.94 (0.05) | 0.98 (0.03) | 0.93 (0.07) | 0.85 (0.11) | 0.98 (0.02) |
|  | **#33** | 0.91 (0.07) | 1.00 (0) | 0.88 (0.10) | 0.79 (0.15) | 1.00 (0) |
|  | **#5** | 0.96 (0.03) | **1.00 (0)** | 0.94 (0.05) | 0.87 (0.09) | **1.00 (0)** |
| **RF** | **#107** | 0.98 (0.01) | 1.00 (0) | 0.97 (0.02) | 0.94 (0.05) | 1.00 (0) |
|  | **#33** | 0.98 (0.01) | 1.00 (0) | 0.97 (0.02) | 0.94 (0.05) | 1.00 (0) |
|  | **#5** | 0.98 (0) | 0.98 (0.03) | 0.98 (0.01) | 0.95 (0.03) | **1.00 (0)** |
| **kNN** | **#107** | 0.83 (0.06) | 0.87 (0.05) | 0.81 (0.08) | 0.65 (0.09) | 0.91 (0.01) |
|  | **#33** | 0.94 (0.05) | 0.98 (0.03) | 0.92 (0.08) | 0.84 (0.12) | 0.98 (0.02) |
|  | **#5** | 0.97 (0.03) | **1.00 (0)** | 0.96 (0.04) | 0.90 (0.08) | 0.99 (0.01) |
| **SVM** | **#107** | 0.96 (0.04) | 0.95 (0.03) | 0.96 (0.04) | 0.91 (0.10) | 0.96 (0.03) |
|  | **#33** | 0.97 (0.02) | 0.97 (0.04) | 0.97 (0.02) | 0.93 (0.06) | 0.97 (0.02) |
|  | **#5** | 0.98 (0.01) | 0.95 (0.03) | **0.99 (0.01)** | **0.98 (0.03)** | 0.97 (0.02) |
| **J48** | **#107** | 0.98 (0.02) | 1.00 (0) | 0.97 (0.02) | 0.93 (0.05) | 0.98 (0.01) |
|  | **#33** | 0.97 (0.02) | 0.97 (0.04) | 0.97 (0.02) | 0.94 (0.05) | 0.99 (0.01) |
|  | **#5** | 0.97 (0.03) | 0.95 (0.03) | 0.97 (0.03) | 0.94 (0.07) | 0.98 (0.03) |

Based on the above results, J48, RF, and SVM have high prediction performances across various metrics, yielding over 0.95 in accuracy, sensitivity, specificity, and AUC on all three levels of features. As a comparison, the state-of-the-art expert algorithm [11] leads to performance of 0.84 in accuracy, 0.78 in sensitivity, 1.00 in specificity, and 0.71 in AUC. This indicates that our features constructed at all the three levels can identify T2DM subjects much better than the popular expert algorithm. State-of-the-art expert algorithm performs slightly better (and almost perfectly) in terms of specificity (1.00) and precision (1.00). This seems most likely due to its stringent conditions on case selection (e.g., a case subject should satisfy any two of the three metrics: diabetic diagnosis, diabetic medications and diabetic lab tests). Obviously, none of our controls satisfy the requirements of cases set above via expert algorithm, and thus bringing the specificity of the expert algorithm to 1 in our experiments.

LR has the highest accuracy (0.99) at the third level of features (as shown in Figure 2 and Table 2), with several other models closely following its performance, such as RF and SVM (with 0.98 in accuracy).

In terms of sensitivity, according to Figure 4 and Table 2, most models experience performance improvement as features get summarized into higher levels. This indicates that simple feature engineering can boost the performance. Meanwhile, multiple models (e.g., LR, NB, and kNN) achieve (near) perfect sensitivity, namely 1.00, on the second or third levels of feature sets. This implies that our framework is highly efficient in make full use of all available data, especially regarding valuable case subjects which are often much rarer and more difficult to accumulate for subsequent studies (such as GWAS and PheWAS).

Accuracy and sensitivity of all classifiers at the third level of features are more stable than on the other two levels of features as shown in Figure 3 and Figure 4, which indicates the summarized final 5 features are stable discriminators to identify T2DM subjects.

For specificity shown in Figure 5 and Table 2, half of the classifiers have performance greater than 0.95 (e.g., RF, SVM, and J48). LR and kNN performance worst when leveraging the first level of features. This may be due to sparsity of features (thus many features end up being noise) or correlated features, which can bias such classifiers.

The AUC performance of our framework also exhibits similar encouraging results. In brief, when trained over second-or third-level feature sets, all classifiers (except LR) manage to perform well above 0.95, which is significantly better than random guessing (0.50) and almost approaching the perfect 1.00. State-of-the-art expert algorithm [11] only scored 0.71 in AUC, making it significantly worse than all models in our framework.

Roughly speaking, as is demonstrated across different metrics in Figures 3 to 7, there is a general trend of increasing predictive performance, as features are abstracted into higher levels. This demonstrates the importance of our feature engineering approach. In addition, we observe better performance improvement from feature engineering than from choices of different machine learning models. This implies that when sample sizes are not sufficiently large (as in most EHR settings), a better strategy to maximize performance should be to refine features.

Overall, across all major metrics, models such as RF, J48 and SVM are more stable than the other three classifiers (kNN, NB, and LR) across the three levels of features. This may be because RF, J48 and SVM are less influenced by sparsity and noise of EHR data, whereas LR, kNN and NB are more vulnerable to these issues.

## Discussion

Traditional expert algorithms use a wide range of separation to select cases and controls, and as a result, a large number of cases and controls are missed. In order to reduce missing rates of current studies, we propose a machine learning-based framework to identify cases and controls in a narrower separation range. We evaluated our framework through Chinese EHR data, and the experimental results show our framework can achieve higher performances than the state-of-the-art algorithm in such EHR data.

However, this work is a pilot study, which is limited in the following aspects:

Firstly, the number of samples (cases and controls) we studied needs to be enlarged in future. Although current selected 221 samples achieve high identification rates on detecting cases and controls, we still need more samples from our repository to confirm scalability of our models. For instance, we can use our classification models to select candidate cases and controls from 23,281 diabetes related patients, and then submit them to clinicians for reviewing. Under such semi-supervised way, we can gain more samples to enrich our framework via a large scale of training (e.g., on more diverse cases and controls) and testing (e.g., on independent new unseen samples). This process will require more reviewing efforts from humans, and will be considered as our next plan.

Second, our framework still involves human efforts in designing of features and confirmations of cases and controls. Although we spent a large amount of time on designing of features, we believe our extracted features could be utilized in other related studies without involving human efforts, which could save them huge amount of time. The evaluations of cases and controls are used to feed our machine learning models. According to achieved high performances of our classification models, researchers can use our model to select cases and controls with a high accuracy, which will save them time to get cases and controls through expert assessments.

Third, compared with expert algorithms in terms of high specificity (small number of non-T2DM are considered as T2DM), our models have lower specificity. This is because we include most of patients between the separation range of cases and controls in expert algorithms (the range between two dotted lines as shown in Figure 2), and as a result, it is hard to make sure all selected cases are predicted correctly. If a study focuses on accuracy of T2DM patient identification more than on number of T2DM patients required, then expert algorithms would be a better choice. If number of cases and controls has higher priority, then our framework would be a better choice.

Fourth, our framework is not confirmed on EHR data of other institutes such as western EHR data. Although the framework achieves a high performance on Chinese EHR data, we believe such EHR data based strategy is also fit for identifying T2DM subjects on western EHR data, and we will test such hypothesis in our next step.

Finally, our methodology focuses on case/control design for traditional association study between phenotypes and genotypes, which requires a perfect precision (wide range of separation between cases and controls in expert algorithms as shown in the Figure 2). The reduced precision rate (leading to a higher recall rates of cases and controls) of our method may influence the traditional association studies. However, as the development of computational phenotyping from EHR data, the association studies will involve more cases with diverse phenotypic characteristics such as comorbidities to enrich the association studies between phenotypes and genotypes. This is because, a disease may be caused by the joint effects of multiple SNPs (i.e. heterogeneity), while a SNP may lead to multiple diseases (i.e. pleiotropy).

## Conclusions

Identifying subjects with and without T2DM is the first step to enable subsequent analysis such as GWAS and PheWAS. In this work, we propose an accurate and efficient framework as a pilot study to identify subjects with and without T2DM from EHR data. Our framework leverages machine learning to automatically extract patterns of T2DM. And we further boost its predictive power by overcoming the wide separation rage of cases and controls in expert algorithms. Our feature engineering framework considers a diverse set of data features spanning diabetic diagnosis codes, diagnosis notes, complications, self-reports, medications (both standard and traditional Chinese medicine), and laboratory tests to represent diabetes related patients. Based on engineered features, we train classification models. We collected 160 T2DM cases and 61 controls and use 4-fold cross validation strategy to evaluate performances of classification models. The experimental results show that our framework can identify subjects with and without T2DM at an average AUC of around 0.98, significantly outperforming the state-of-the-art at an AUC of 0.71.

## Appendices

Appendix A: A list of 110 constructed features

Appendix B: A list of diabetic medicine

Appendix C: A list of positive and negative diagnosis notes related with T2DM

Appendix D: A list of 36 features summarized from 110 features as listed in Table A1

Appendix E: A list of 8 features summarized from 36 features as listed in Table A4

Appendix F: Expert algorithm for the identification of subjects with T2DM

## Acknowledgements

We thank members from the Changning regional distributed EHR network, Shanghai, China, for sharing of de-identified EHR data studied in this paper. We would like to thank physicians from hospitals and medical centers belonging to Changning district to evaluate cases and controls.

## Appendix A A list of 110 constructed features

The constructed 110 features across seven sources are listed in Table A1. For every source, we design specific features covering diagnosis codes (ICD-10 codes E11. ***), diagnosis notes (positive notes and negative notes as shown in Table A3 of Appendix C), self-report notes (persistent hunger, polyuria, and polydipsia), medications (traditional Chinese medicine and western medicine), plasma glucose test (venous and peripheral) and HbA1C test.

**Table A1.**
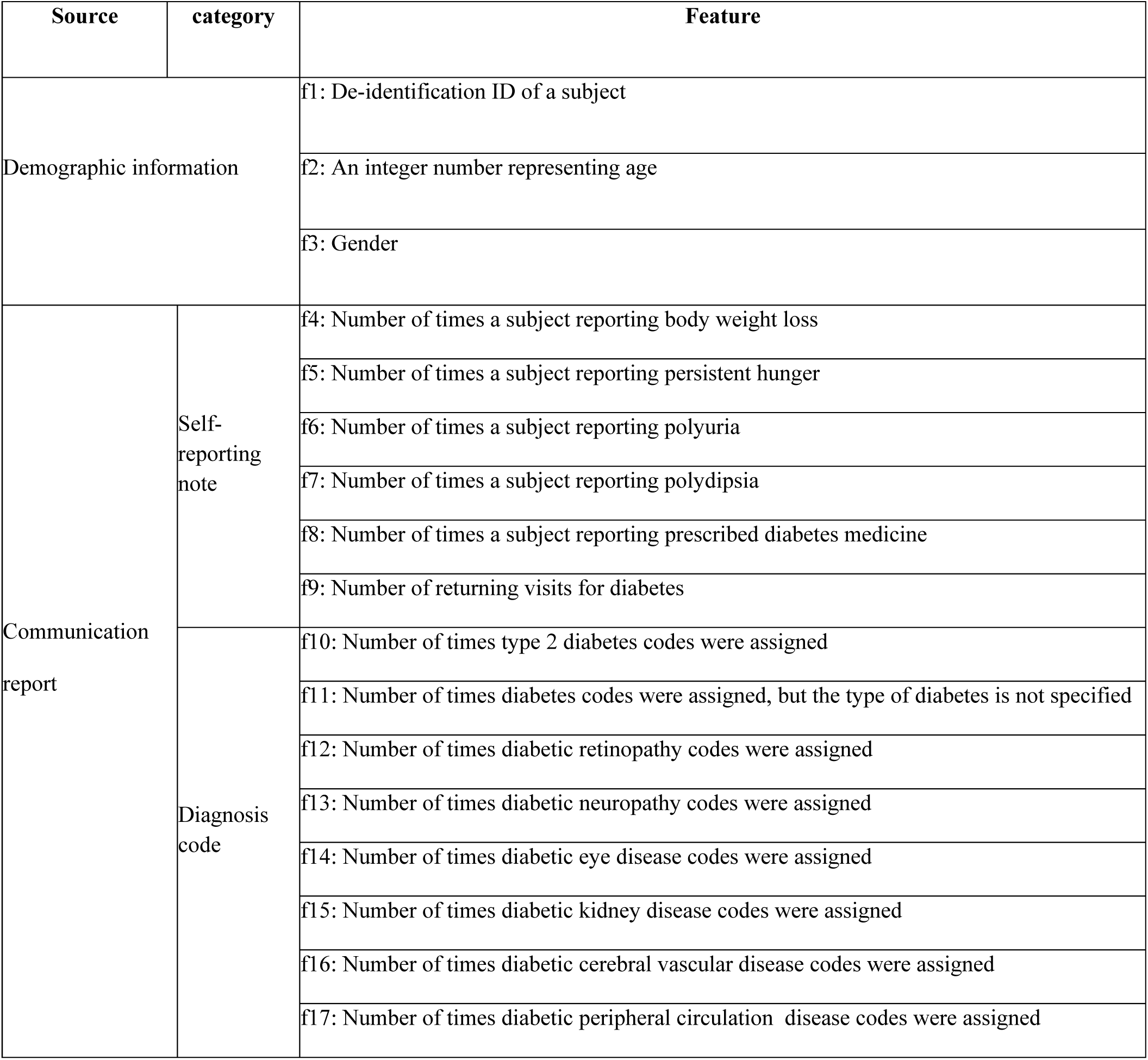
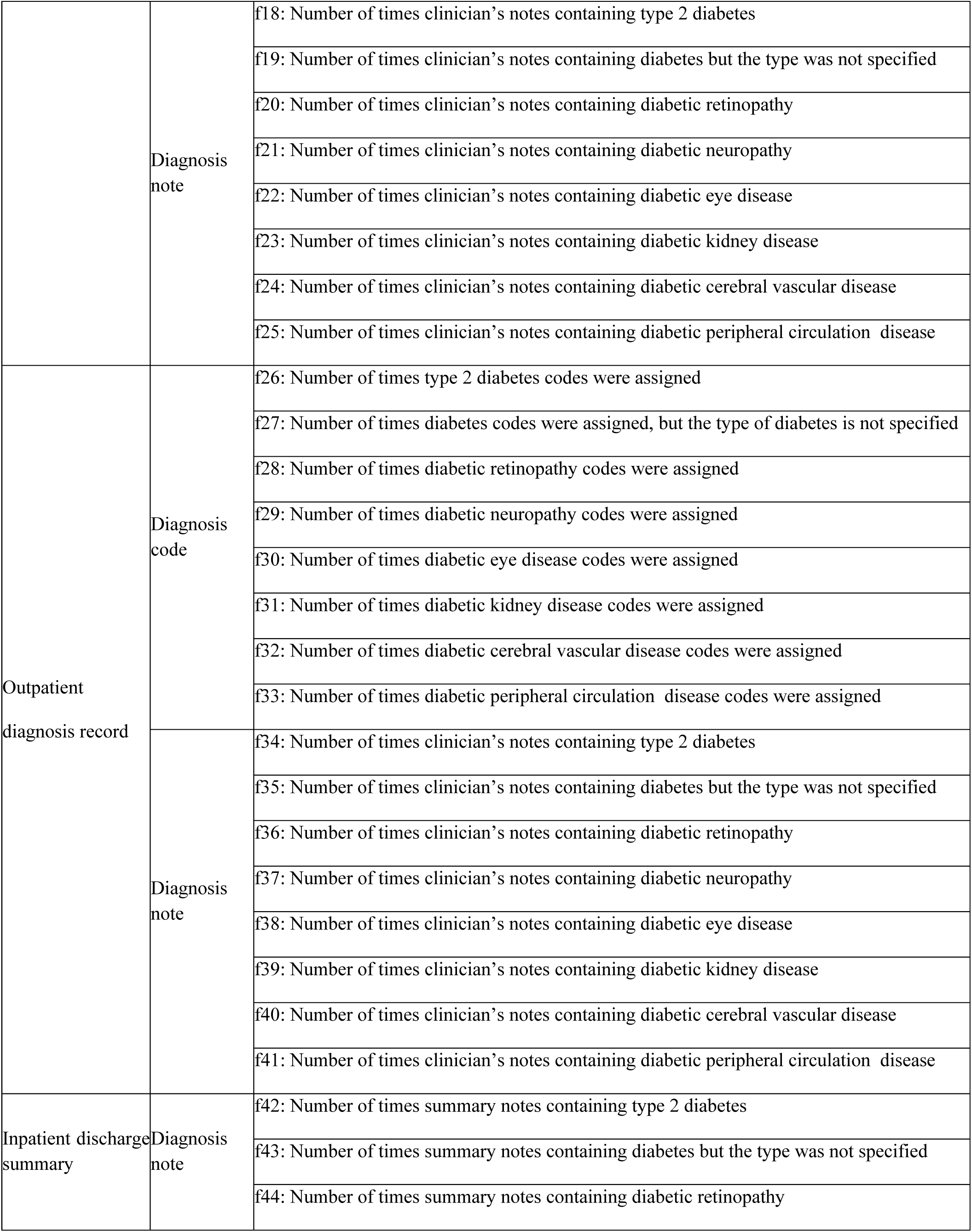
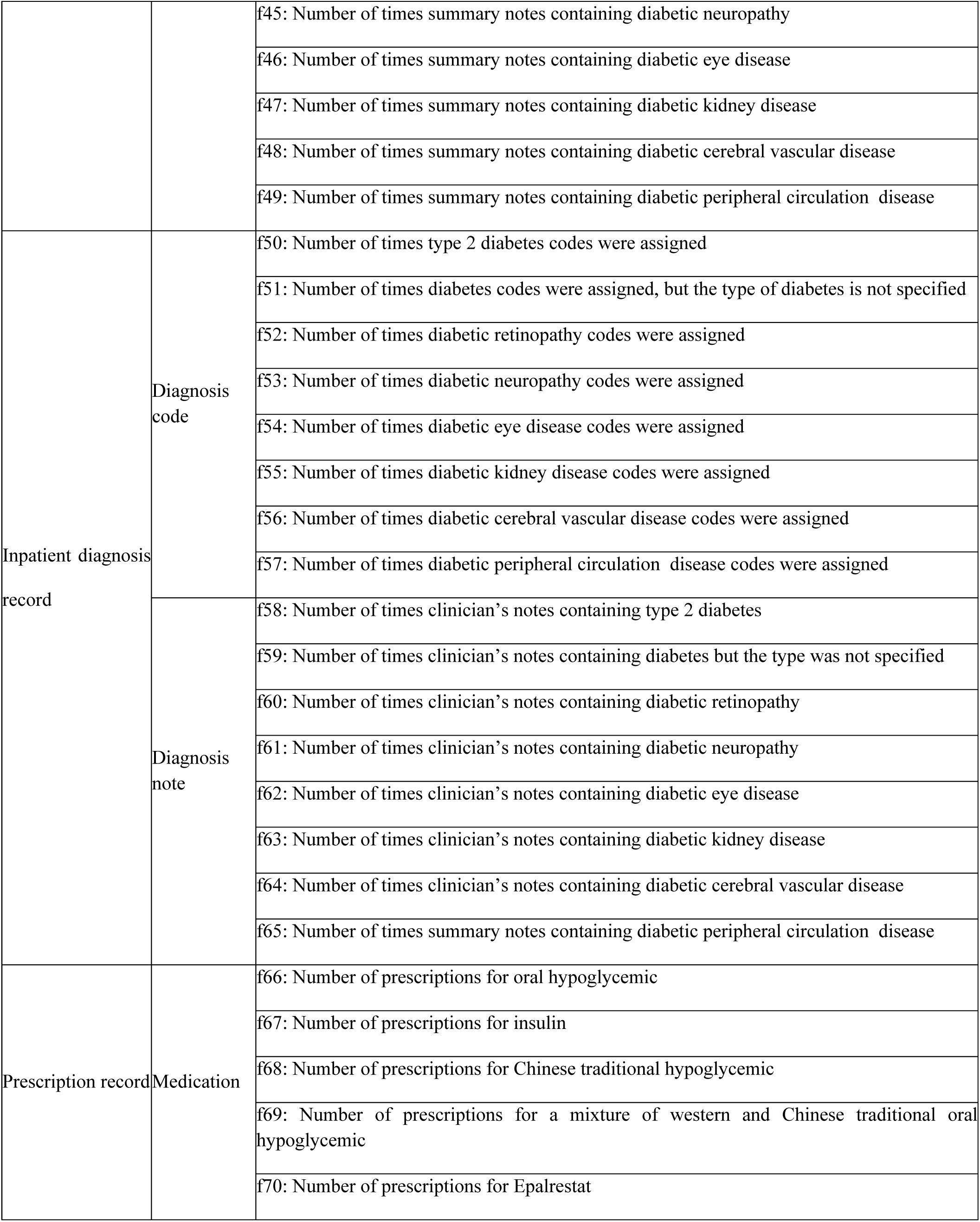
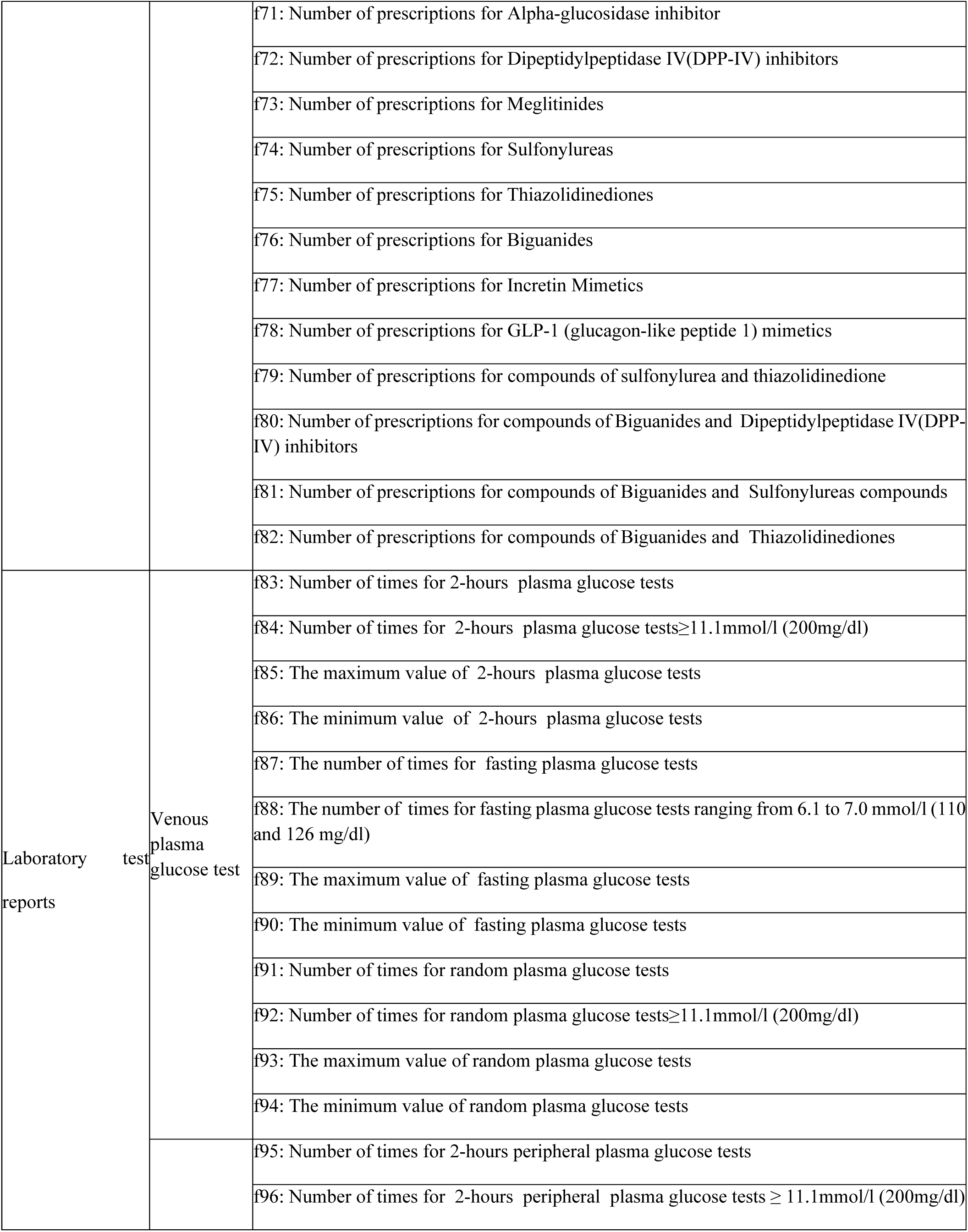
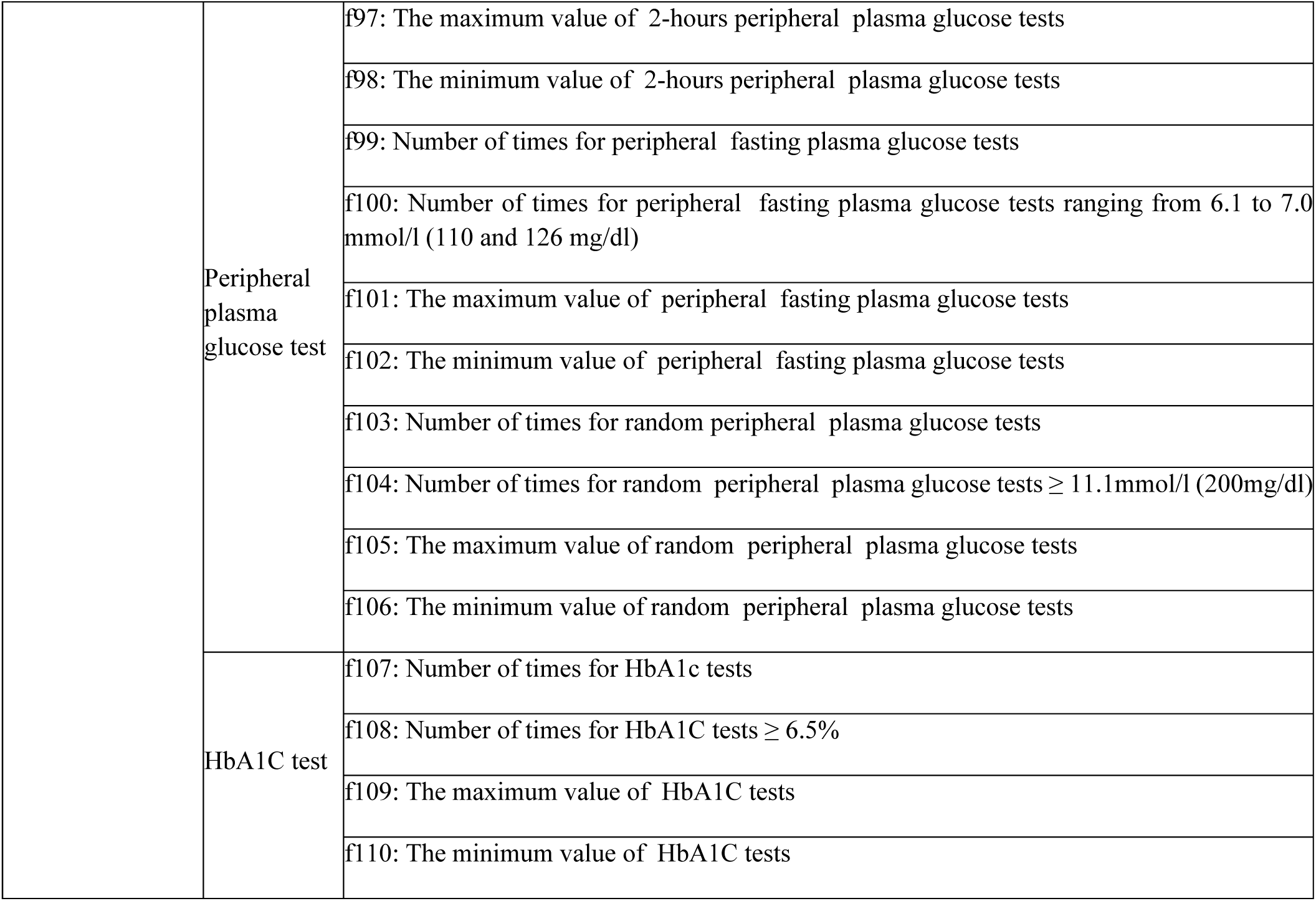
The constructed 110 features coming from seven sources.

## Appendix B A list of diabetic medicine

Medicine is a principal factor to characterize phenotypes of subjects with type 2 diabetes mellitus (T2DM). In this paper, we use prescribed medicine listed in Table A2 as one of our seven sources to construct medicine related features as listed in Table A1.

**Table A2.**
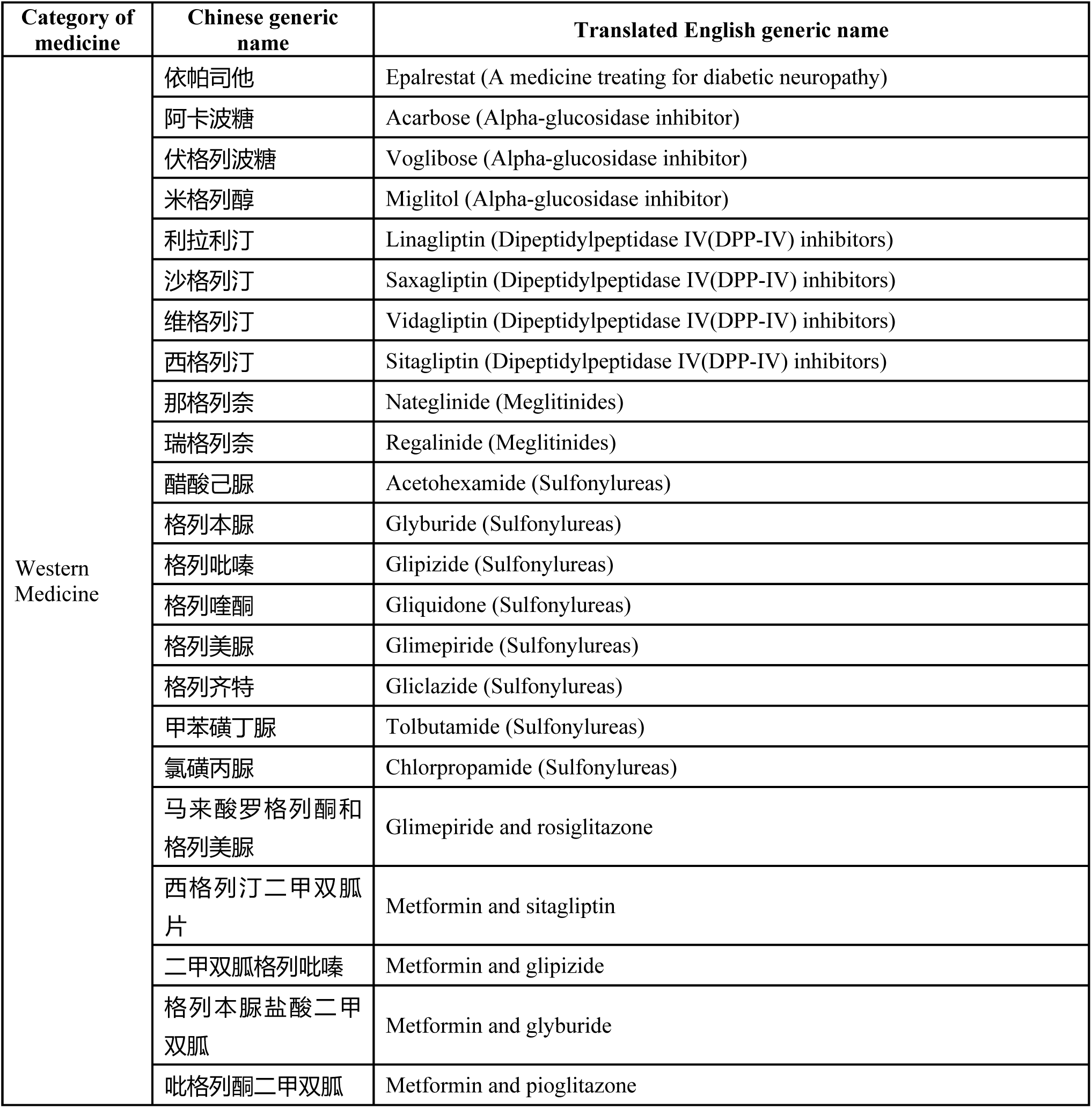
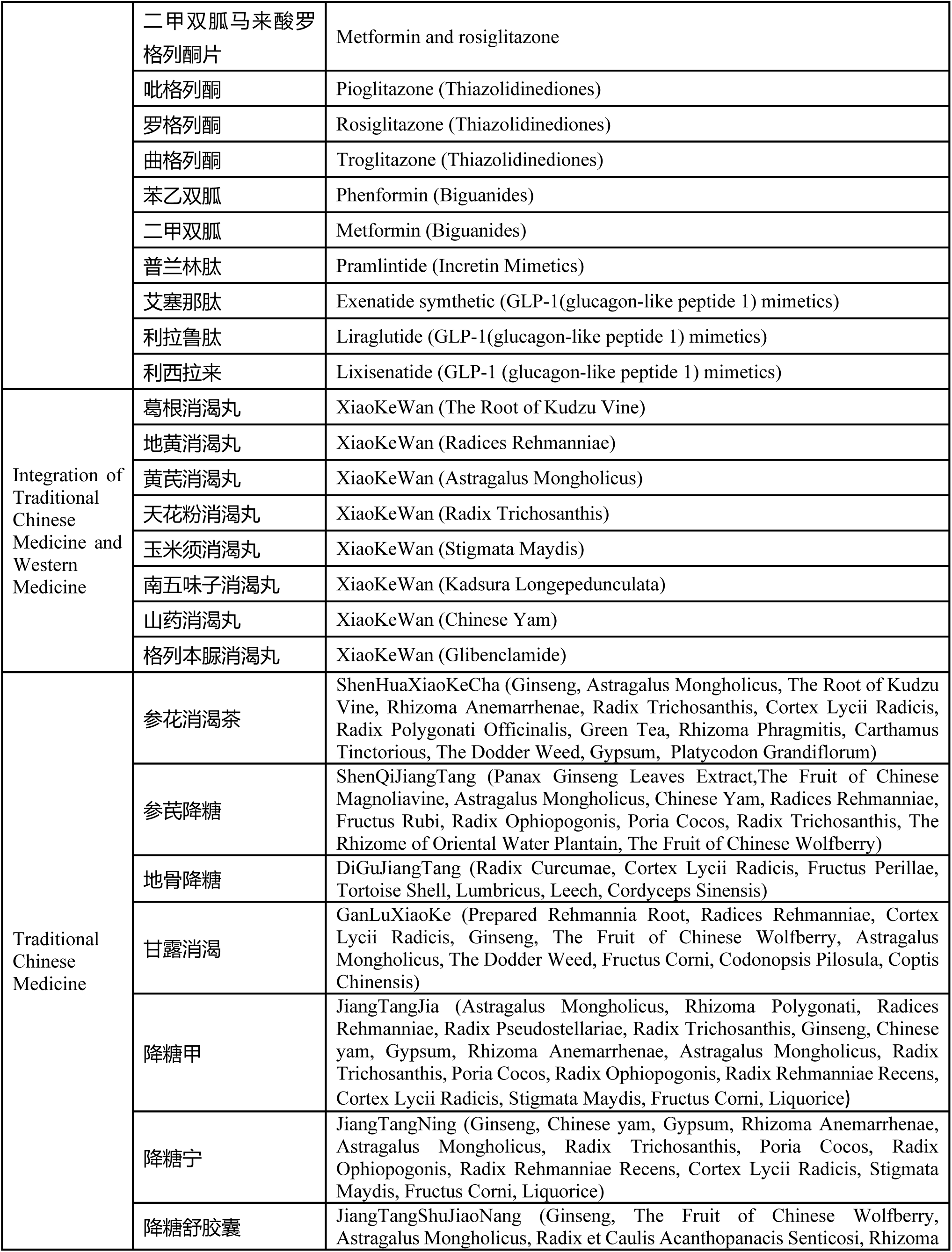
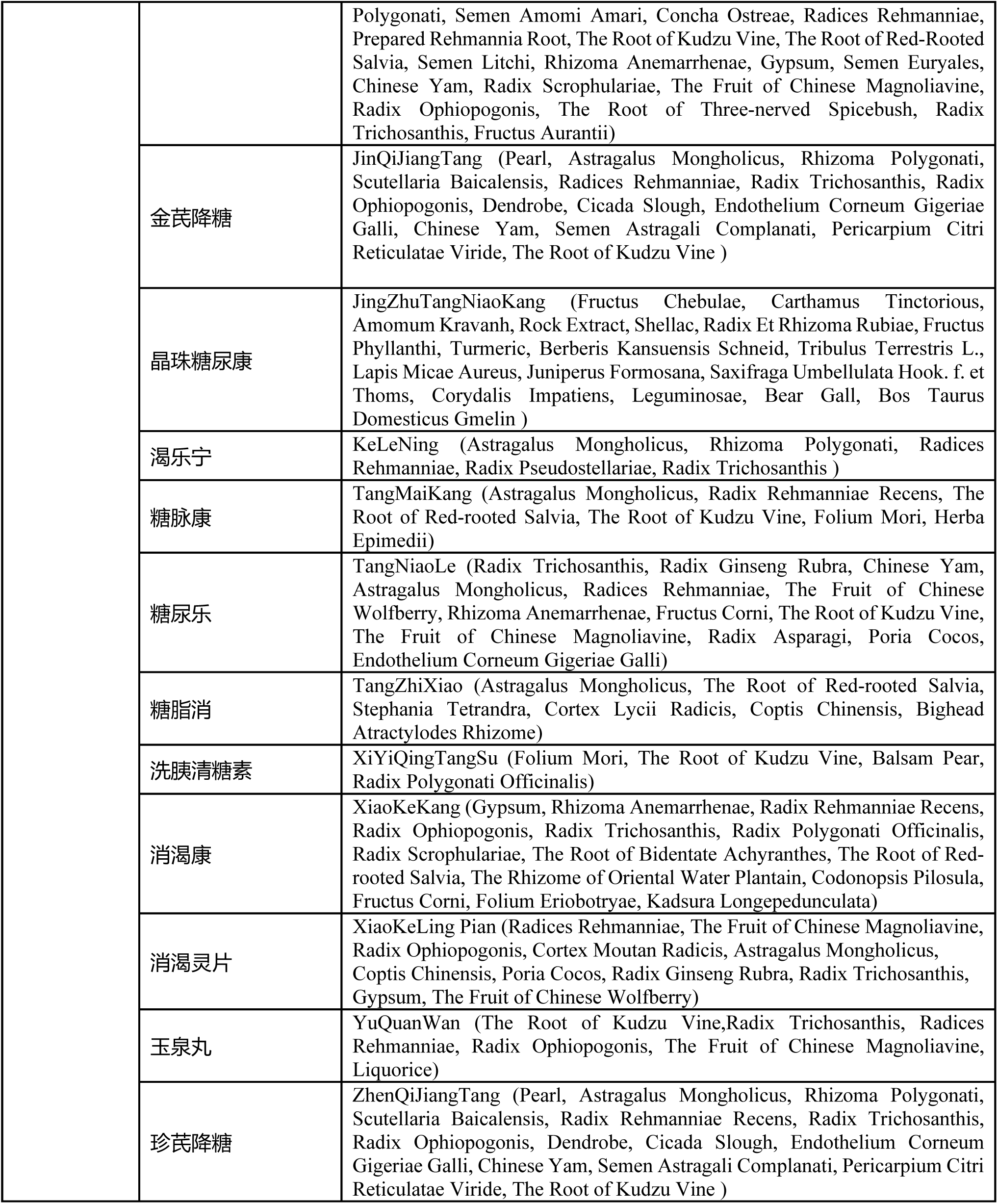
A list of medicine associated with subjects with type 2 diabetes mellitus

## Appendix C A list of positive and negative diagnosis notes related with T2DM

Diagnosis notes existing in diagnosis reports or clinical summaries are represented as unstructured texts. We create a dictionary of diagnosis notes related with T2DM. There are two types of diagnosis notes: positive and negative. We assume that if a subject’s EHR data contains positive diagnosis notes, but not negative diagnosis notes, then the positive diagnosis notes are counted to construct features associated with diagnosis notes.

**Table A3.**
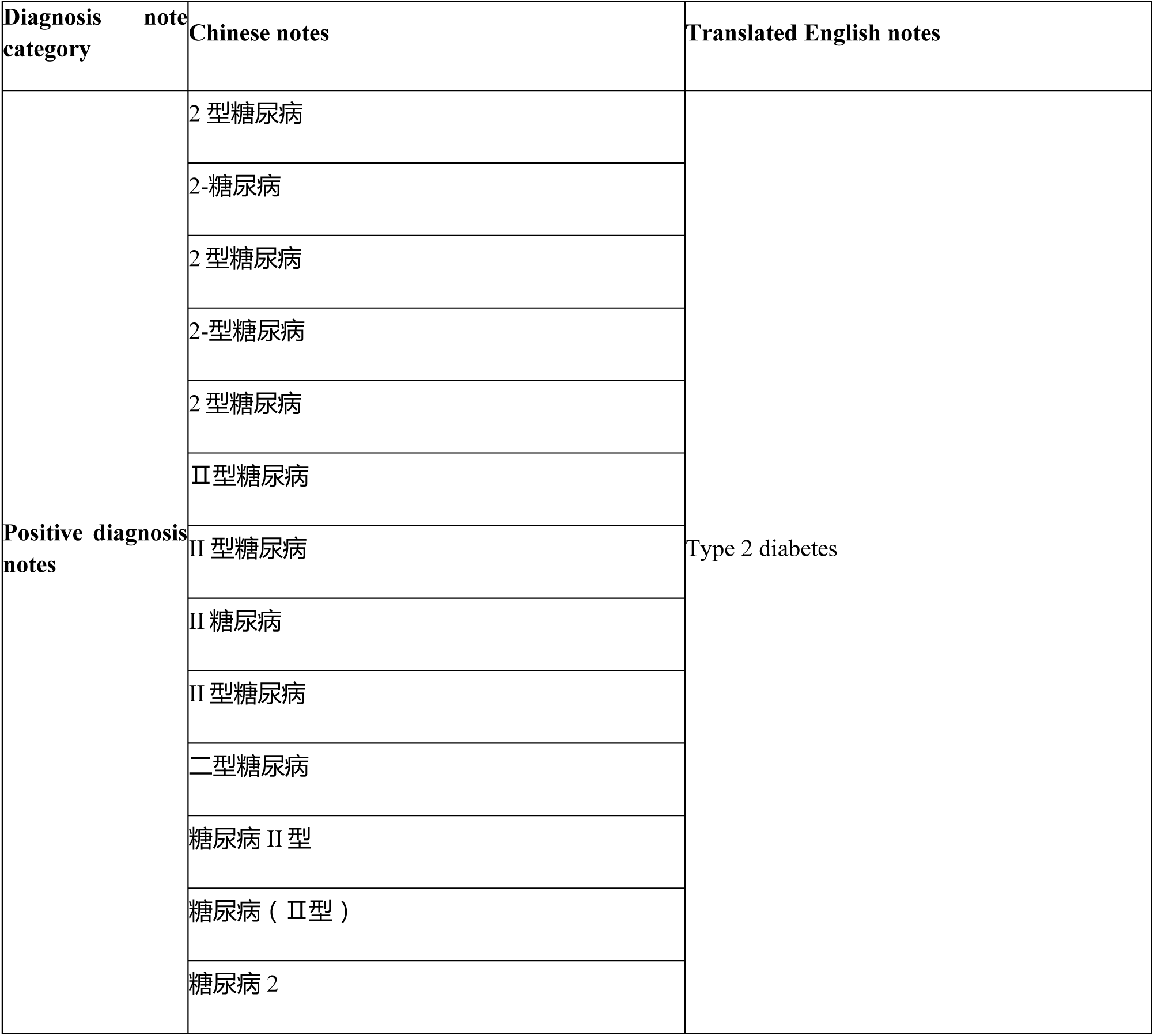
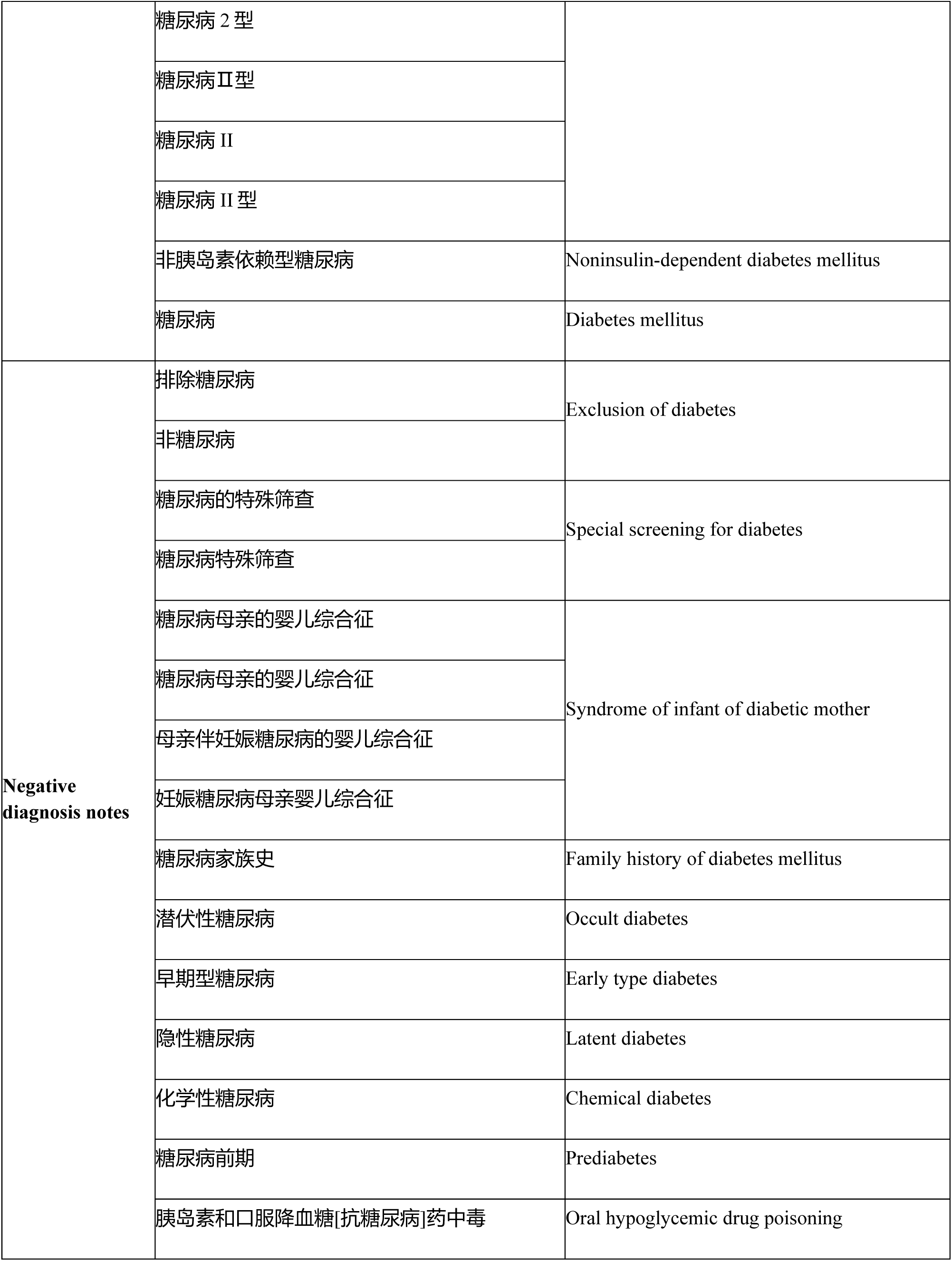
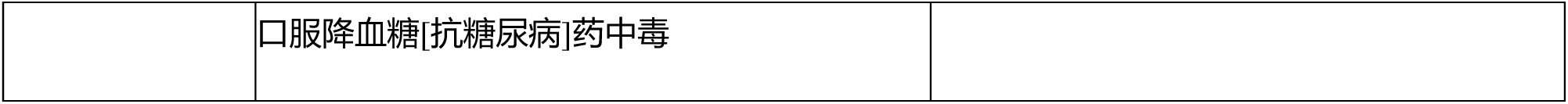
A list of medicine associated with subjects with type 2 diabetes mellitus

## Appendix D A list of 36 features summarized from 110 features as listed in Table A1

Features listed in Table A1 are extracted from seven sources, however, several features across sources are correlated. For instance, diagnosis-code related features appearing in “*communication report*”, “*outpatient diagnosis record*” and “*inpatient diagnosis record*” are similar. These features have the same definition in above three sources, so they can be summarized as a new feature. In this way, eight new features (f’_10_ to f’_17_) in the category of diagnosis codes as shown in Table A4 are summarized from 24 features (f_10_ to f_17_, f_26_ to f_33_, f_50_ to f_57_) from Table A1. By using the same way, we summarize 32 similar diagnosis-note related features appearing in “*communication report*” (f_18_ to f_25_), “*outpatient diagnosis record*” (f_34_ to f_41_), “*inpatient diagnosis record*” (f_42_ to f_49_) and “*inpatient discharge summary*” (f_58_ to f_65_) into 8 new features (f’_18_ to f’_25_) in the category of diagnosis notes as shown in Table A4.

Features as listed in the “*laboratory test report*” of Table A1 are also correlated with each other. For instance, features ranging from f_83_ to f_86_ are all correlated with venous 2-hours plasma glucose test. In order to reduce negative influences of correlated features on the performances of classification models such as k nearest neighbors, we only keep features which are positive signals of type 2 diabetes. For instance, feature f84 characterizing the number of times 2-hours plasma glucose test≥11.1mmol/l, which is a positive signal of type 2 diabetes conditions. So do feature f_88_, f_92_, f_96_, f_100_, f_104_ and f_108_.

>Most of subjects only take a small number of medicine listed in Table A2, as a result, the data covering features ranging from f_66_ to f_82_ has a big sparsity, which will influence the performances of computational models to learn patterns of T2DM [1]. In order to avoid a big sparsity, we transform original features ranging from f_66_ to f_69_ into new ones ranging from f’_26_ to f’_29_ as shown in Table A4.

**Table A4.**
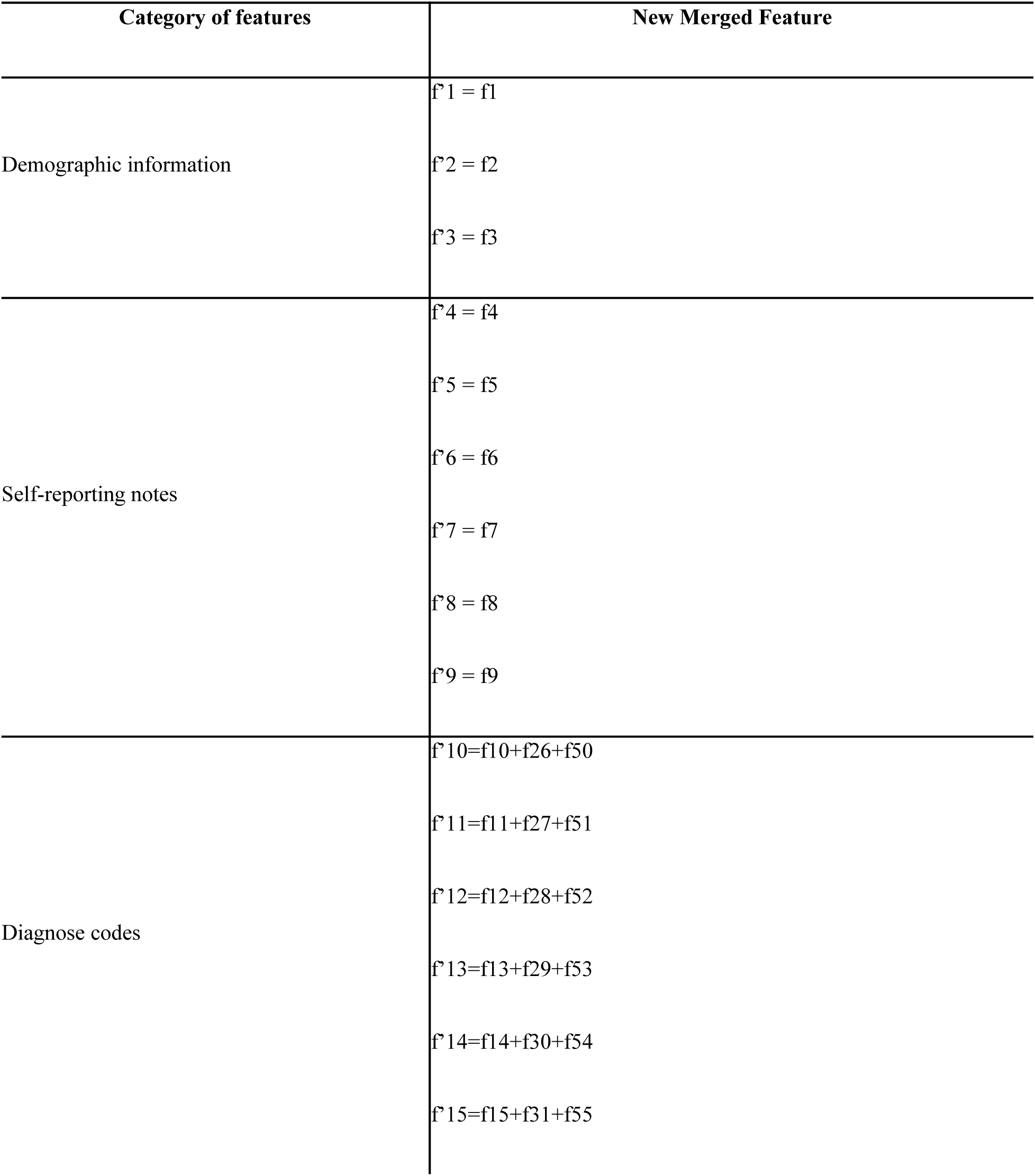
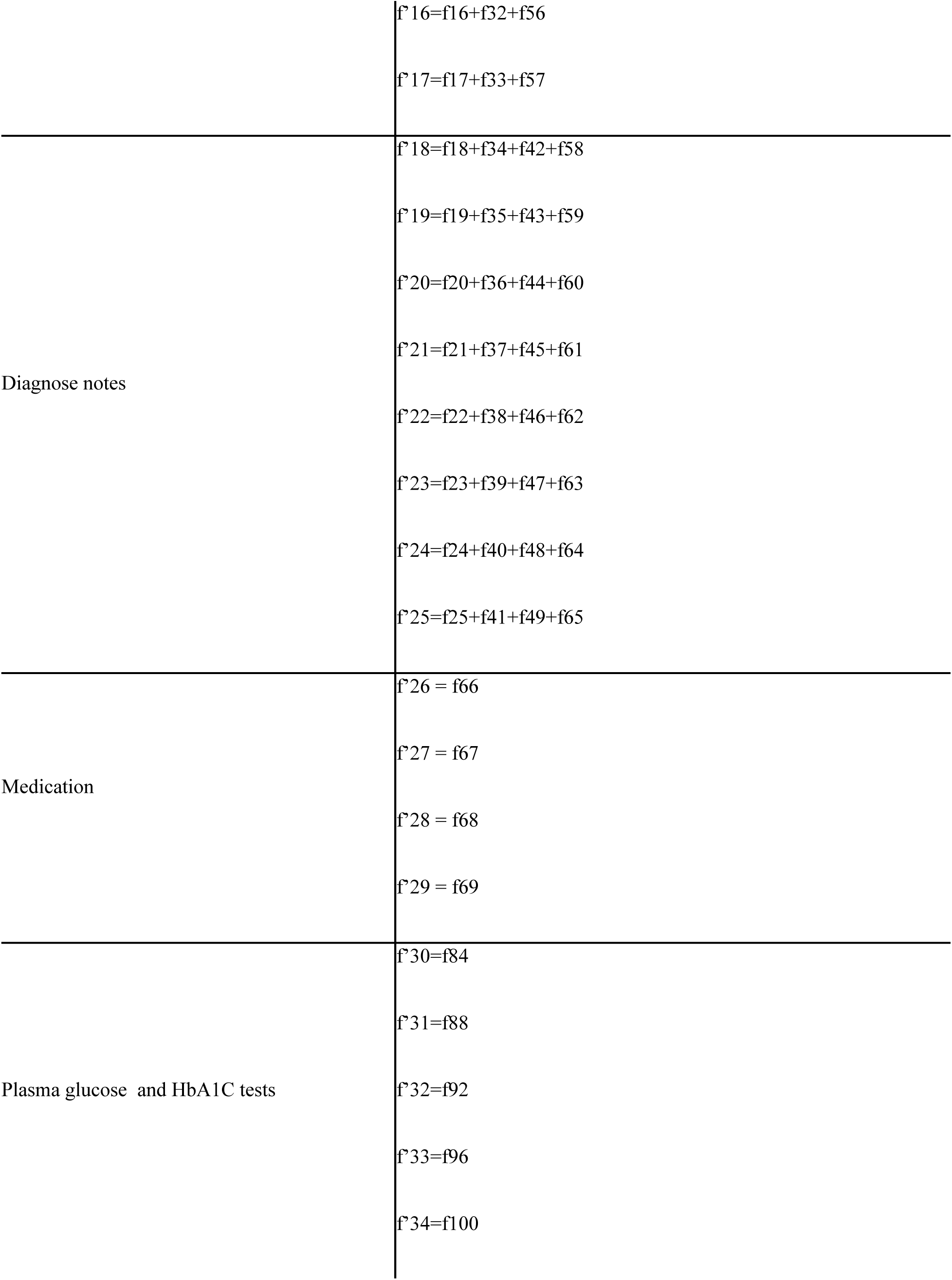
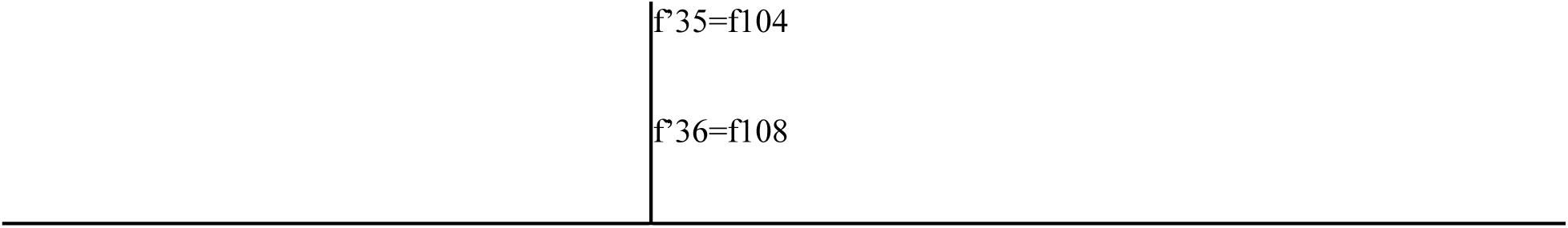
The original 110 constructed features as shown in Table A1 are transformed into 36 features via summarizing similar features across seven sources: “*communication report*”, “*outpatient diagnosis record*”, “*inpatient diagnosis record*”, “*inpatient discharge summary*”, “*prescription report*” and “*laboratory report*”.

## Appendix E A list of 8 features summarized from 36 features as listed in Table A4

36 features in Table A4 are summarized as 8 features in following 6 categories:

1. Patients’ demographic information ranging from f’’_1_ to f’’_3_.
2. Self-report summarize 6 features ranging from f’_4_ to f’_9_ in Table A4 as f’’4 in Table A5s to represent the total number of times diabetic phenomena such as body weight loss, persistent hunger, polyuria, polydipsia, prescribed diabetes medicine and returning visits for diabetes were reported by subjects in the source of “*communication report*”.
3. Diagnosis code summarize 8 features ranging from f’_10_ to f’_17_ in Table A4 as f’’_5_ in Table A5 to represent the total number of times diabetic diagnosis-codes are assigned to a subject in “*communication report*”, “*outpatient diagnosis record*” and “*inpatient diagnosis report*”.
4. Diagnosis note summarize 8 features ranging from f’_18_ to f’_25_ in Table A4 as f’’_6_ in Table A5 to represent the total number of times diabetic diagnosis-notes are described in a subject’s “*communication report*”, “*outpatient diagnosis record*”, “*inpatient diagnosis record*” and “*inpatient discharge summary*”.
5. Medication summarize 4 features ranging from f’_26_ to f’_29_ in Table A4 as f’’_7_ in Table A5 to represent the total number of times diabetic medicines as listed in Table A2 are prescribed in a subject’s prescription record.
6. Plasma glucose and HbA1C test summarize 7 features ranging from f’_30_ to f’_36_ in Table A4 as f’’8 in Table A5 to represent the total number of times venous plasma glucose, peripheral plasma glucose (fasting plasma glucose ≥126 mg/dl or 2-hours plasma glucose≥200 mg/dl or random plasma glucose ≥200 mg/dl) and HbA1C tests are abnormal.

**Table A5.**
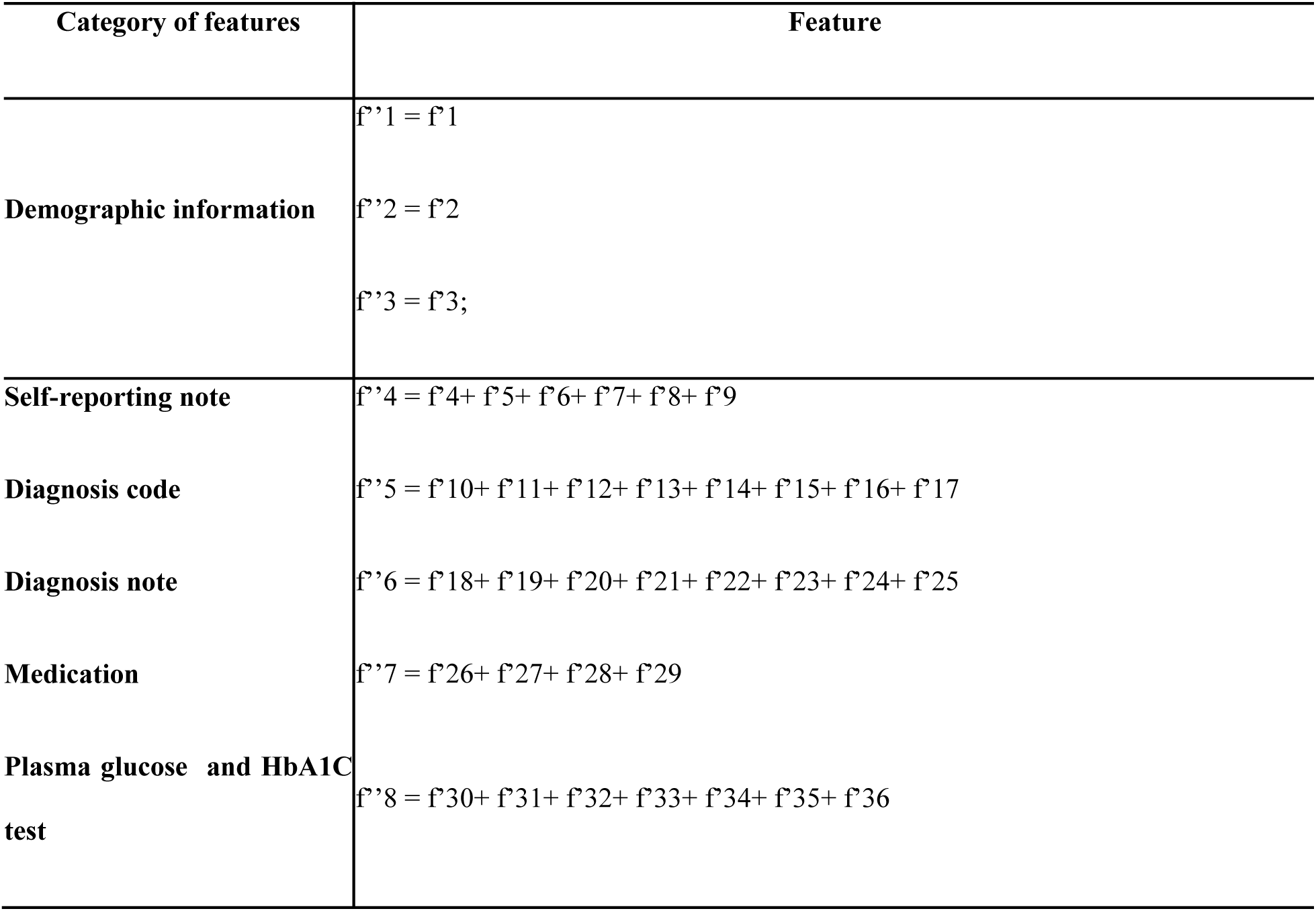
The 8 features after summarizing related features within a category such as “self-reporting note”, “diagnosis code”, “diagnosis note”, “medication”, “plasma glucose” and “HbA1C test”.

## Appendix F Expert algorithm for the identification of subjects with T2DM

The expert algorithm^2^ we used as our baseline to do performance comparisons is depicted in Figure A1. The performance of the algorithm had been successfully validated at multiple eMERGE Network^3^ sites in the USA. The algorithms utilized various types of information including diagnosis codes, medication orders, laboratory results and clinical notes. We applied this algorithm on all of our investigated EHR sources including diagnoses, laboratory results, medications, communication reports and clinical notes. Notably, the expert algorithm and our approach both used the same EHR sources.

**Figure A1.**
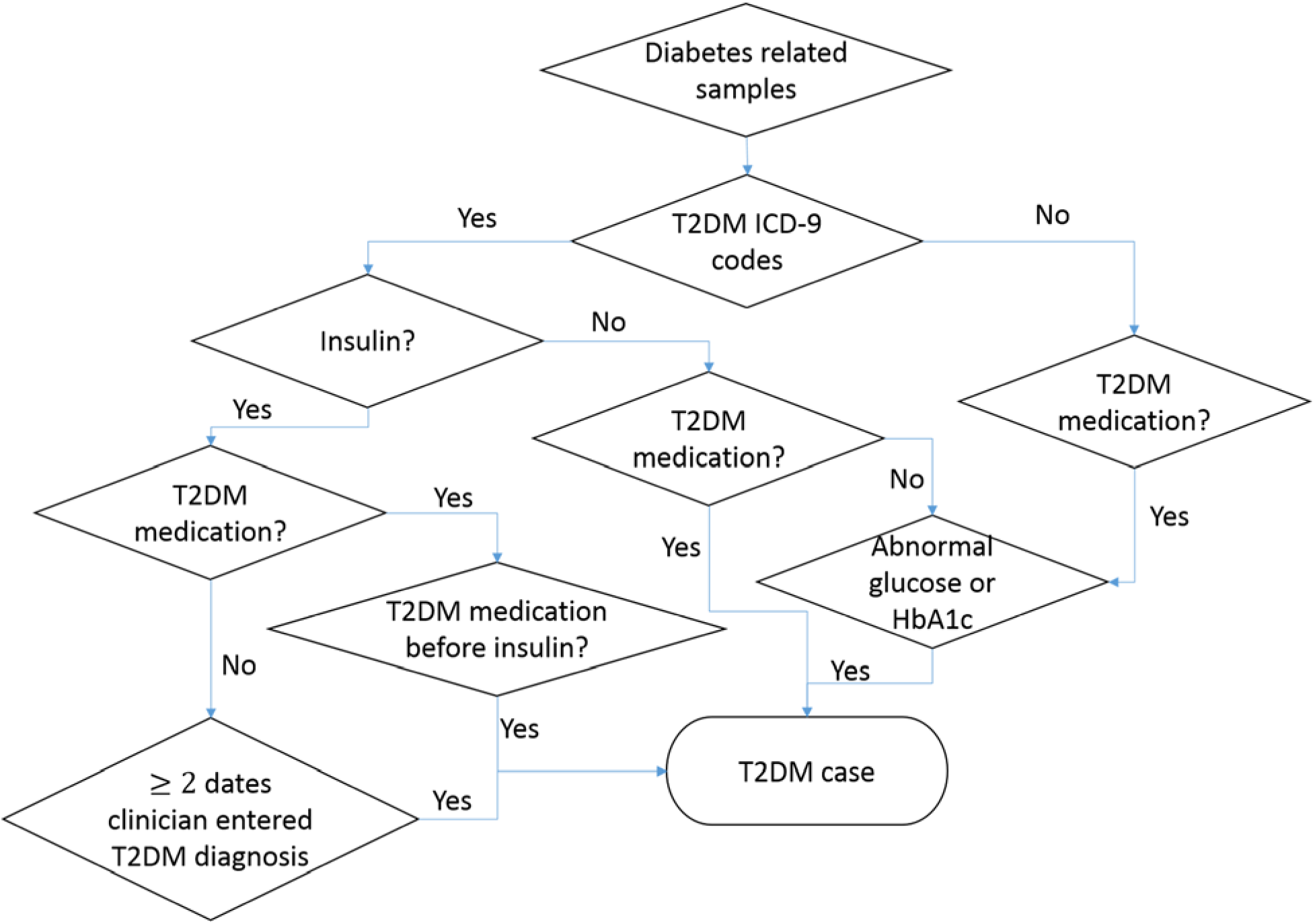
Expert algorithm for the identification of subjects with T2DM

## Footnotes

1 We exclude three demographic features, because they do not indicate any significant differences between cases and controls, and in contrast, they will influence the correct determinations of classifiers such as kNN.

